# A reference dataset of 5.4 million phased human variants validated by genetic inheritance from sequencing a three-generation 17-member pedigree

**DOI:** 10.1101/055541

**Authors:** Michael A. Eberle, Epameinondas Fritzilas, Peter Krusche, Morten Källberg, Benjamin L. Moore, Mitchell A. Bekritsky, Zamin Iqbal, Han-Yu Chuang, Sean J. Humphray, Aaron L. Halpern, Semyon Kruglyak, Elliott H. Margulies, Gil McVean, David R. Bentley

## Abstract

Improvement of variant calling in next-generation sequence data requires a comprehensive, genome-wide catalogue of high-confidence variants called in a set of genomes for use as a benchmark. We generated deep, whole-genome sequence data of seventeen individuals in a three-generation pedigree and called variants in each genome using a range of currently available algorithms. We used haplotype transmission information to create a phased “platinum” variant catalogue of 4.7 million single nucleotide variants (SNVs) plus 0.7 million small (1-50bp) insertions and deletions (indels) that are consistent with the pattern of inheritance in the parents and eleven children of this pedigree. Platinum genotypes are highly concordant with the current catalogue of the National Institute of Standards and Technology for both SNVs (>99.99%) and indels (99.92%), and add a validated truth catalogue that has 26% more SNVs and 45% more indels. Analysis of 334,652 SNVs that were consistent between informatics pipelines yet inconsistent with haplotype transmission (“non-platinum”) revealed that the majority of these variants are *de novo* and cell-line mutations or reside within previously unidentified duplications and deletions. The reference materials from this study are a resource for objective assessment of the accuracy of variant calls throughout genomes.

## Introduction

Recent disruptive changes in sequencing technology (Bentley et al. 2008; Drmanac et al. 2010) have led to a massive growth in the use of DNA sequencing in research and clinical applications (The 1000 Genomes Project Consortium 2010; The International Cancer Genome Consortium 2010; Erikson et al. 2016). Accurate calling of genetic variants in sequence data is essential as sequencing moves into new settings such as clinical laboratories (Gullapalli et al. 2012; Goldfeder et al. 2016). It is anticipated that genomic sequence information will improve the precision of clinical diagnosis as part of the new initiatives in precision medicine (Ashley 2015; Marx 2015). The field of next-generation sequencing (NGS) is evolving rapidly: continual improvements in technology and informatics underline the need for effective ways to measure the quality of sequence data and variant calls, so that it is possible to perform objective comparisons of different methods. Robust benchmarking enables us to better understand the accuracy of sequence data, to identify underlying causes of error, and to quantify the improvements obtained from algorithmic developments.

It is important to assess aspects of variant calling accuracy such as the fraction of true variants detected (recall) and the fraction of the variants called that are true (precision). One approach is to test variant calls made by an NGS method using an orthogonal technology (e.g. array-based genotyping or Sanger sequencing) and then to measure the degree of concordance between results (Ajay et al. 2011; The 1000 Genomes Project Consortium 2012; Pirooznia et al. 2014). This approach can provide a measure of precision of a variant caller, but not recall, as recall estimates require knowledge of what is missed. Additionally, the resulting measure of precision is typically based on a few hundred variants, and is then extrapolated to the entire variant call set. Limitations in this approach to validation include cost, and incompleteness due to failed or erroneous results from the orthogonal technology. A second approach is to compare technical and/or informatic replicates of a dataset (Lam et al. 2012; O’Rawe et al. 2013; Zook et al. 2014). It is assumed that a variant call is correct if it is seen in multiple analyses or datasets. While this approach allows rapid comparison of large variant call sets, estimates of precision and recall of one variant call set can only be expressed relative to a second set; and it is not possible to know which variants are true in either set. Additionally, calls found in two datasets may be categorized as correct even where they are in fact systematic errors in both sets. A significant limitation of this approach is that some of the variants called by just one method may be correct and may provide valuable insights on how to improve variant calling, but these variants are excluded from further consideration by this approach. A third approach is to sequence parent-parent-child trios and test for Mendelian consistency (Boland et al. 2013; Patel et al. 2014). While this approach can detect a subset of errors it falls short of identifying genotyping errors that do not violate inheritance in a trio (see supplementary methods).

In the present study, we generated a genome-wide catalogue of 5.4 million phased “platinum” variants. We included variant calls from six different informatics pipelines (Conrad et al. 2011; Garrison and Marth 2012; Iqbal et al. 2012; Saunders et al. 2012; Raczy et al. 2013; Rimmer et al. 2014) and two different sequencing technologies (Bentley et al. 2008; Drmanac et al. 2010). We used an inheritance-based validation based on a family of two parents and 11 children to resolve conflicts between different call sets, and to include high confidence variants called by a single informatics pipeline. Compared to previous studies, the genetic inheritance provided by this large family serves as a validation of all of the variants within this study and provides an unbiased assessment of different technologies and software pipelines. We also examined features of the SNVs that failed to segregate consistently with the inheritance and conclude that the majority of these conflicts are caused by SNVs that co-localize with copy number variants (CNVs). The data from this study are fully open-source so that groups can reanalyze and provide feedback to the catalogue as needed. As the technologies improve, this community-resource will continue to evolve by adding more validated variants including CNVs and structural variants (SVs) identified from new sequencing technologies and analysis methods.

## Results

### Phasing the pedigree

To obtain phased variants across a large pedigree we sequenced each of the four grandparents, two parents and eleven children of CEPH pedigree 1463 (Dausset et al. 1990) (Figure 1) on an Illumina HiSeq2000 to an average depth of 50x using 2x100 bp reads and PCR-free sample preparation. We determined the genome-wide transmission of the parental haplotypes to each of the 11 children in the pedigree and identified 731 inheritance vectors between parents and children within the autosomes, plus 16 distinct inheritance vectors on chromosome X (see Methods). In agreement with previous studies (Kong et al. 2010), we observed a higher number of crossovers in maternal versus paternal autosomal haplotypes: 58% (415/709, 1.35 cM/Mb) compared to 42% (302/709, 0.98 cM/Mb) (supplementary methods).

**Figure 1.**
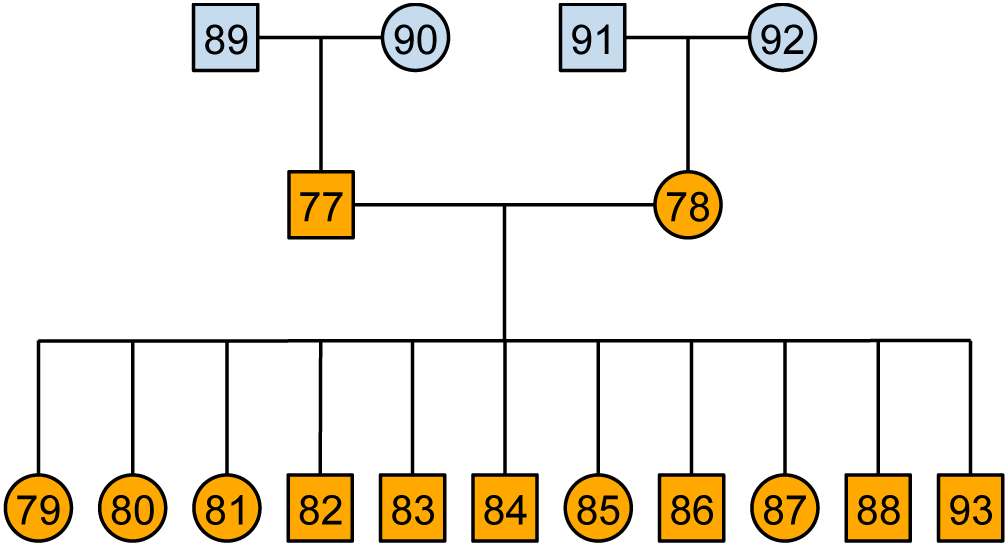
Pedigree of the family sequenced for this study (CEPH pedigree 1463). The Coriell ID for each sample is defined by adding the prefix NA128 to each numbered individual: e.g. 77 = NA12877. Samples filled with dark orange are used in this analysis but the founder generation (blue) were also sequenced and used as further validation of the haplotypes generated in this study. The trio, 91-92-78 was also sequenced during Phase I of the 1000 Genomes Project (The 1000 Genomes Project Consortium 2010).

Sequencing and phasing a larger pedigree increases the ability to detect errors and assess the accuracy of more of the variants compared to a trio analysis. Theoretically, by sequencing the parents and 11 children, both parental haplotypes will be represented in at least one of the children for 99.8% of the genome, and single errors in variant calls can be detected in all of these regions in any member of the pedigree for which sequence data are available (supplementary methods & Table S6). To this effect, combining the genotype calls with the haplotype inheritance vectors enables identification of genotype errors as well as other factors that lead to Mendelian inheritance inconsistency such as when variants co-segregate with deletions and duplications.

### Platinum variants

To generate a comprehensive catalogue of pedigree-validated variants we employed six analysis pipelines (supplementary methods) to call SNVs and indels. For each individual call set we identified variants where the genotypes are present in all thirteen individuals and are also consistent with the transmission of the parental haplotypes. Next we merged each of the six call sets into one catalogue, using the following inclusion criteria: (a) the genotypes and alleles were concordant between call sets (154,681 variants removed by this rule) and (b) for monoallelic variants (i.e. homozygous for the alternate allele in every sample), the variant was included in call sets using at least two different sequence aligners (103,603 variants removed by this rule). Note that we included all variants that met the above criteria regardless of whether or not they passed the default quality filters of each analysis pipeline. Instead, we relied on the sensitivity of the genetic inheritance to detect genotyping errors and maximize the chance of including true variants that might otherwise be removed by suboptimal filtering (see supplementary methods).

It is still possible that misalignments may lead to false genotypes that are consistent with the inheritance or a single variant may be represented as two different, non-overlapping variants when merging the call sets. To identify errors such as these, we required that some of the reads supporting each variant call also supported the flanking sequence using a k-mer approach. In brief, we declared each variant valid only if the supporting reads contained sequence confirming each allele and the flanking sequence in a 51 bp window centered on the variant (see supplementary methods). We performed this check for every variant in the catalogue and removed 132,528 SNVs and 64,917 indels where the genotype calls were consistent with inheritance but failed this additional flanking-sequence test.

In total, we identified 5,426,236 phased variants (4,729,676 SNVs, 693,623 indels and 2,937 complex variants with overlapping SNVs and indels) that segregate consistently with haplotype inheritance and also pass the flanking-sequence test including 159,167 pedigree-consistent variants that were identified from a single sequencing pipeline (see supplementary methods). These variants have features consistent with high quality sequencing data. For example, the ratio of transitions to transversions in the pedigree-validated SNVs is 2.11 (all) and 2.42 (coding) and the het:hom ratios of SNVs and indels in both the parents of this family are in agreement with both previous studies and theoretical predictions (also see supplementary methods). In addition to the 2,937 complex variants, the set also includes 59,310 multi-allelic variants (2,728 SNVs, and 56,582 indels). The indel loci include short tandem repeats such as homopolymer tracts, di- and tri-nucleotide repeats (see supplementary methods) which are known to be less stable than SNVs, hence we expect a significant number of the indels to be multi-allelic, as observed here.

To identify invariant positions of high confidence we collated all genomic positions that were called as homozygous reference in every sample by at least two pipelines. As for the monoallelic variants (described above) we required that the different pipelines included at least two different sequence aligners. Additionally, we filtered out all positions where any pipeline made a variant call in any sample to exclude missed variants and regions of high error rates. Based on these rules we identified 2,737,246,156 positions that are homozygous reference across the pedigree. These positions can be used to calculate false positive rates when assessing variant calling pipelines.

### Using founder haplotypes to validate platinum variants

We utilized the founders to provide an additional assessment of the phasing and variant quality by testing whether the predicted haplotype variants are observed in the appropriate founders (i.e. individuals 89, 90, 91 or 92 in Figure 1). All the platinum variants identified by this study are phased, and one of the haplotypes will be present in each founder. To quantify the presence of the variant in the founders without relying on the variant callers, we tested if the k-mers described above were observed in the appropriate founders. This analysis supported 99.5% of the 5,426,236 variants in all of the founders leaving 28,371 lower-confidence variants (supplementary methods). Variants with a low normalized k-mer value are more likely to fail this test due to either lower average depth or higher error rate. For example, ∼97.3% of the Platinum variants have at least 5 k-mers supporting each allele (Table S7) compared with just 16.4% (4,652) of the variants that failed the founder test. After excluding the 23,719 variants with fewer than 5 k-mers supporting each allele, many of the remaining variants exhibit signs of clustering around biological mutational events in a grandparental haplotype. We presume, therefore that they arose either in early development, or during generation or culture of the cell line: specifically 2,066 (46.7%) occur in 135 clusters of between 5 and 260 spatially adjacent variants in the same founder (see examples in Figure 2).

**Figure 2:**
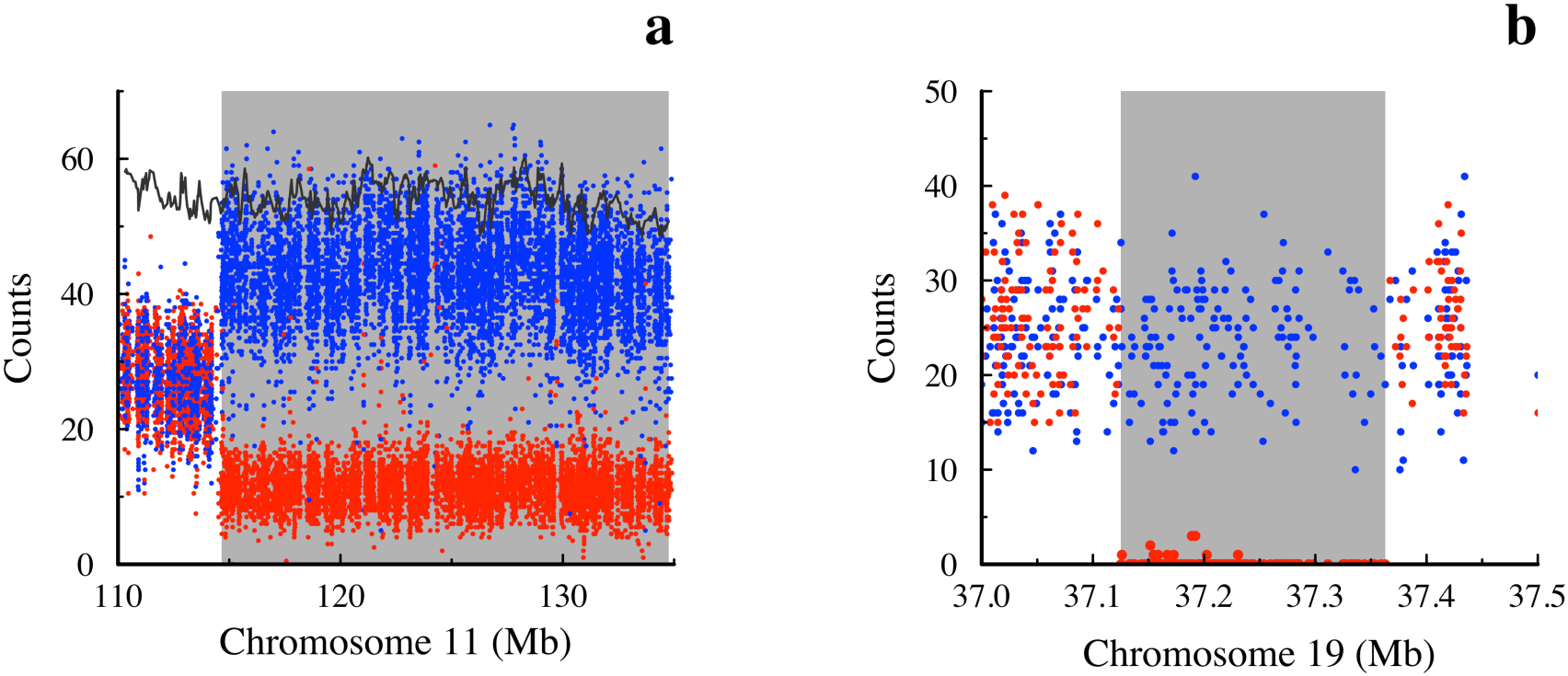
Structural abnormalities account for most of the inconsistencies in detection of the platinum haplotypes in the founders. **(a):** A 20 Mb structural abnormality identified by 177 variants in 9 clusters that failed the founder haplotype validation test in NA12889. Based on the marked skew in frequency of the SNV alleles within the rearrangement (4:1) compared to the proximal flanking region (1:1), this event is likely a mosaic on the distal end of chromosome 11, specific to NA12889. Each point represents the allele count at a SNV location expected to be heterozygous in NA12889 based on seeing the non-transmitted allele at least 6 times. Red points show the allele counts (average n=11 within the mosaic) of the haplotype transmitted to NA12877 and blue points show the allele counts (average n=43 in the mosaic) of the non-transmitted haplotype. The black line shows the average total depth in windows of 100 SNVs highlighting that this mosaic is not associated with a change in copy number. **(b)**: Allele counts for SNV positions in a possible cell-line somatic deletion in NA12891 identified by a cluster of 174 k-mer failures. Points are colored the same as in (a). Within this deletion, there are virtually no reads corresponding to the transmitted haplotype, while a relatively constant read depth was observed across the region for the non-transmitted haplotype.

It should be noted that these biological mutational events are limited to the founders and thus do not affect the pedigree consistency or our assessment that these variants are accurately called in the main pedigree. Overall, for the variants where at least one of the founders did not have the appropriate kmer, 99.2% show strong supporting evidence for the variant either because the k-mer was seen as predicted in another founder (72.2%) and/or the k-mer was seen multiple (5+) times in the pedigree (99.3%). Because of this, we conclude that most of the failures in the founders are caused by complex genetic events in the grandparents and do not affect the concordance of transmission. These variants are flagged but not removed from the Platinum catalogue.

### Non-platinum variants

The non-platinum variants that failed our validation process revealed important biological insights. We examined a total of 334,652 high-quality SNV calls, where at least two pipelines provided the same answer for all genotype calls in the parents and all children, but the genotypes were inconsistent with haplotype inheritance. This analysis excluded all non-platinum SNV calls that might be of low quality, e.g. where there were missing data, or discrepancies between two callers, or variants called by only one caller. We classified the high quality failed SNVs into four categories as follows (see supplementary methods for detailed analysis): Category 1 SNVs (191,087) are heterozygous in all thirteen individuals. Most (91%) of the SNVs in this category are clustered at specific locations in the genome, and also deviate significantly (p<0.01) from Hardy Weinberg Equilibrium (HWE) in a population of European ancestry (unpublished data), having an excess of heterozygous genotypes. Together these observations indicate that most of these variants overlap real duplications or higher order CNVs, and are not false positive variants (see Figure 3a for an example of a 25kb duplication). Category 2 SNVs (3,861) are consistent with the occurrence of underlying hemizygous deletions in the family. Most (83%) of the SNVs are clustered (see example in Figure 3b) and have low read depth in samples predicted to be hemizygous for the deletion (25.6x compared to the 50.3x diploid genome average). Many (81%) are also significantly out of HWE in the European cohort. Category 3 SNVs (49,800) are positions with a single heterozygous call in the pedigree. These singletons are not clustered; they may be isolated false positive calls or somatic mutations. Of these, 48.8% (24,299) were called identically in two independent datasets generated by different sequencing chemistries and analysis pipelines (Complete Genomics and Illumina), suggesting that they are likely to be mostly true somatic mutations, arising either in the individual or during culture of the cell line, plus an expected ∼50-100 *de novo* germline mutations in each of the children. The remaining 51.2% (25,501) are just identified in one sequencing technology and there is no evidence to indicate how many are errors versus low frequency somatic mutations. Category 4 SNVs (89,904) are the remaining positions that are not pedigree consistent. Compared to the previous categories, these may be accounted for by a variety of reasons. While some SNVs are genotyping errors, most (79%) of the common SNVs are significantly out of HWE and approximately half (54%) of all the SNVs are clustered, which is consistent with the occurrence of both somatic deletions and *de novo* germline duplications (but does not meet the criteria for inclusion in categories 1-3). An example of a complex event is shown in Figure 3c, where somatic instability in different individuals has occurred in 22q11.2, a region in the genome that is known to be highly mutable (Mikhail et al. 2014). Additionally, there are approximately 589 SNVs in 322 clusters that failed to match the inheritance vectors consistent with a double crossover or gene conversion (see supplementary methods).

**Figure 3.**
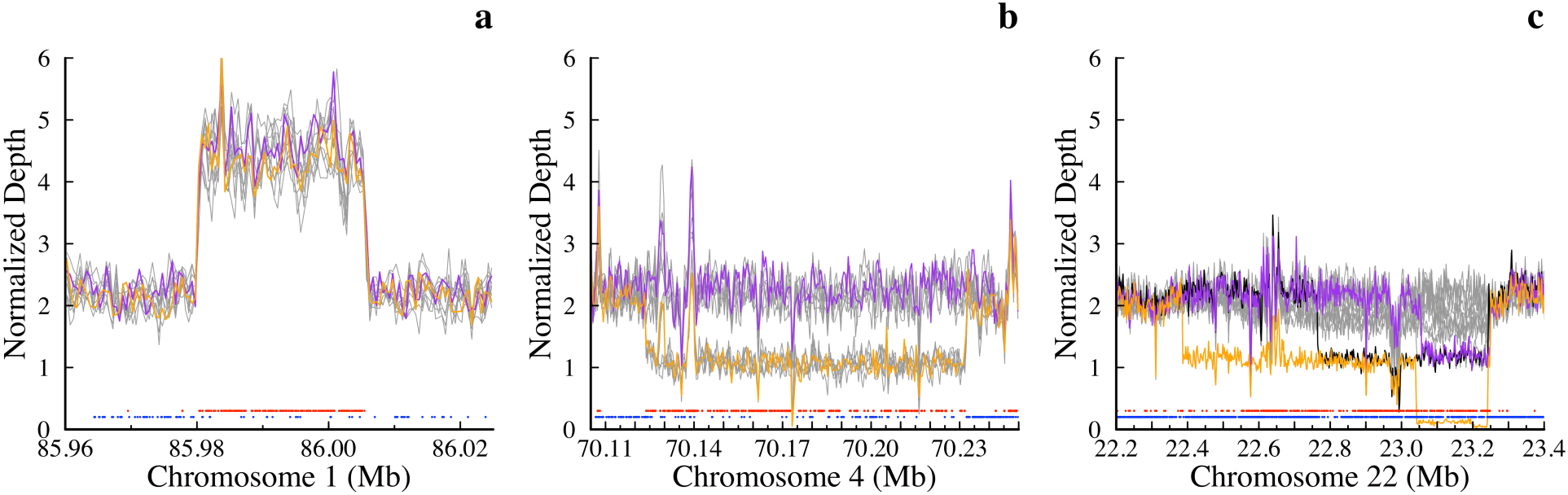
CNVs in this pedigree identified from non-platinum variants. **(a)** Duplication on chromosome 1 containing 242 category 1 non-platinum SNVs in a region of elevated read depth. Colored lines show the depths for the parents (NA12877 in purple & NA12878 in orange) and the grey lines show the depths for each of the children. Points along the bottom highlight the platinum variants (blue) and the non-platinum variants (red). Note that there are 16 platinum variants in this duplication because the presence of duplicated sequence can still produce genotypes that are consistent with those predicted by a diploid model (see supplementary methods). **(b)** Deletion on chromosome 4 identified by 176 category 2 non-platinum SNVs that were consistent with the presence of a large hemizygous deletion. In addition to the large deletion, the depth supports the presence of several segmental duplications both overlapping and flanking the deletion. Lines and points are colored as above. **(c)** Cell line or somatic deletions in multiple members of the pedigree on chromosome 22 identified by 926 category 4 non-platinum SNVs. Lines are colored as above except for the addition of a black line that shows the depth for NA12893 (child). While the other children (grey lines) do not appear to be deleted in any part of this region the depths are highly variable, which may indicate somatic instability and mosaicism. The variability of this region within different cell line passages is evident when we compared this sequence data for NA12878 against corresponding data from the 1000 Genomes Project and NIST (see supplementary methods).

Comparing our non-platinum SNVs with a set of common, population-level CNVs (Sudmant et al. 2015a) and CNV calls from this pedigree (Roller et al. 2016) shows a significant excess of Category 1 and Category 4 SNVs overlapping duplications and significantly more Category 2 SNVs overlapping deletions (see supplementary methods). Overall, we conclude that the majority (∼75%) of non-platinum SNVs in this analysis co-locate with an unidentified germline or somatic variant. 67.3% were accounted for in 23,442 clusters (indicative of CNVs). Improvements in CNV calling and modifying the inheritance analysis to utilize the copy number information will lead to more accurate characterization of these events. While many of these are likely to be true variant locations, assessing algorithm performance should exclude these sites until improved CNV calling is available and variants pass the necessary criteria for inclusion in the platinum variant catalogue (see Discussion).

### Using the Platinum variant catalogue for benchmarking

We used the Platinum variant catalogue to benchmark the performance of four commonly used informatics pipelines on sample NA12878, using WGS at 30x, 40x or 50x average aligned read depths (Figure 4 and supplementary methods). We performed joint calling with the parental datasets for those callers that have a multi-sample analysis mode. For precision, we utilized the homozygous reference positions identified as high confidence. For SNVs, FreeBayes has the highest recall at the cost of lower precision compared to the other algorithms. The results for Strelka and GATK3 are similar, with GATK3 recall showing slightly higher recall and lower precision than Strelka. For indels, Strelka and GATK3 have very similar recall and precision, while Platypus has the highest precision values at a cost of lower recall. We used default filters for these comparisons, but recall can be improved for any algorithm with a resulting loss in precision by altering filter thresholds (see for example the receiver operator characteristic (ROC) curves for GATK3 and Strelka in Figure 4b).

**Figure 4.**
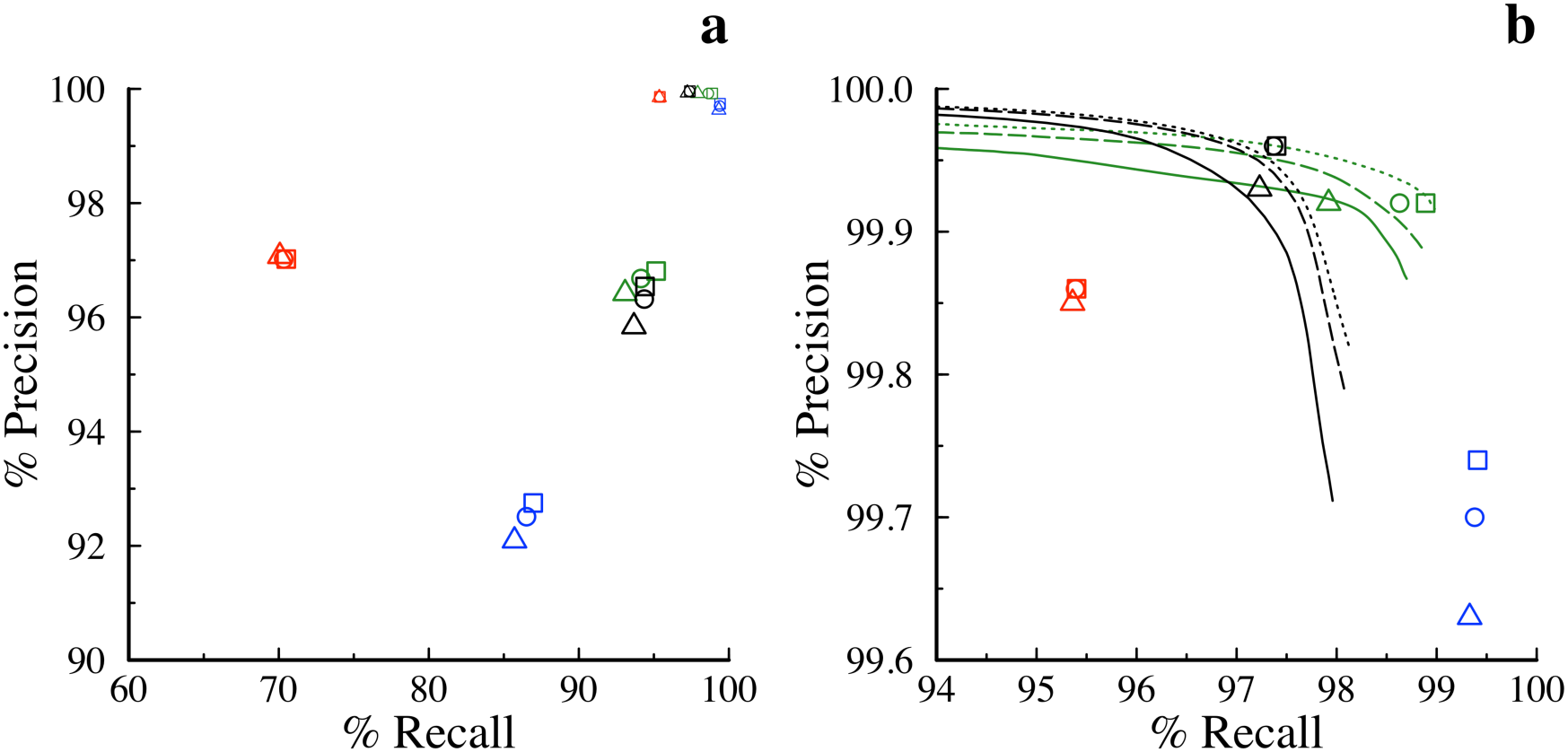
Precision versus recall in NA12878 evaluated against the Platinum catalogue dataset. Triangles, circles and squares respectively represent the results from 30x, 40x and 50x sequencing depth for Platypus (red), FreeBayes (blue), GATK3 (green) and Strelka (black). Excluding Strelka, all callers are run in joint calling mode incorporating the parents. **(a)** Indels (large symbols) and SNV (small symbols) results plotted on the same axis. **(b)** Expansion of SNV results, also showing ROC curves for GATK3 and Strelka that reflect the trade-off of recall versus precision that is obtained by altering specific variable parameters when using the algorithms.

### Comparison to other studies

The NIST Genome-in-a-Bottle (Zook et al. 2014) and the 1000 Genomes (1KG) (The 1000 Genomes Project Consortium 2015) both included data from NA12878 and we compared variant calls from these studies with our Platinum variant catalogue. All three studies generated high-confidence, genome-wide variant sets and then used different approaches to filter the calls and assess quality. The goal of the NIST study was to generate a high-confidence set of variants as a resource for benchmarking SNV and indel calls, and is therefore directly aligned with the aim of the present study. The goal of the 1000 genomes project was to generate a comprehensive catalogue of human genetic variation across a diverse set of individuals from multiple populations across the globe.

The NIST study was a replicate-based analysis of a single sample (NA12878) using 14 separate sequencing experiments (total depth = 831x) (Zook et al. 2014). Variants were excluded from the final set by arbitration based on inconsistent genotype calls between pipelines, and/or occurrence in repetitive/duplicated sequence or structural variants. In contrast to our Platinum genome calls, NIST used neither pedigree information nor flanking-sequence analysis to identify errors, though visual inspection was used for some variants. We found that the genotype concordance was very high for the SNVs (>99.99%) and indels (99.92%) called in both datasets (Table 2). The Platinum dataset also contained 801k SNVs and 224k indels not included in the NIST set. Additionally we identified 912,018 SNVs and 261,647 indels that are more difficult (based on the observation that they were not called consistently by all of the pipelines) including 347,186 (38.2%) SNVs and 142,964 (54.6%) indels not included in the NIST set (see supplementary methods). Identifying these “difficult” variants is particularly important because they highlight areas where the variant calling methods can be improved. Given that these additional variants passed our inheritance and flanking-sequence tests, these can be considered true positives. Most of these variants (91.3% of the SNVs and 63.5% of the indels) were also found independently in the 1KG dataset (see below). Conversely, the NIST dataset contained ∼63k SNVs and 60k indels absent from the Platinum dataset. The majority of these were observed in this study but were then filtered out (see supplementary methods).

**Table 2.**
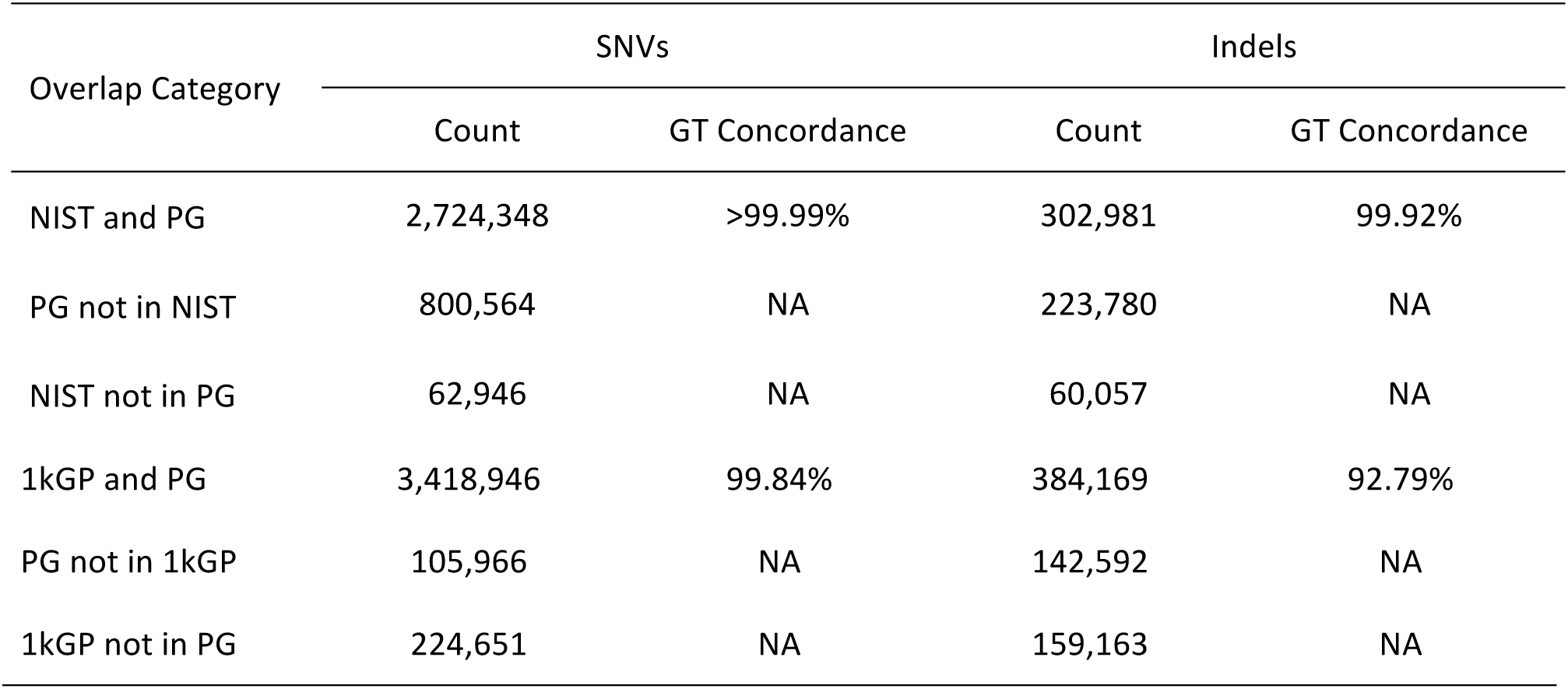
Comparison of Platinum genotype calls in NA12878 with NIST and 1KG.

The 1KG dataset was generated from 2,504 individuals (including NA12878) using 7.4x average depth WGS data combined with 65.7x exome sequence data and array-based genotype data (The 1000 Genomes Project Consortium 2015). Haplotype-based imputation was used for error-correction and to impute missing data. This cohort-based analysis was calibrated to maintain a population-level false discovery rate (FDR) of <5% for most variants, including SNVs and small indels. Genotyping concordance for the 3.4M SNVs shared between the two datasets is high (>99.8%), but lower (92.8%) for the indels. The Platinum dataset contains 105,966 SNVs and 142,592 indels that are absent from the 1KG data for NA12878. When we examined the entire 1KG dataset, we found 28% of these SNVs and 30% of the indels were in fact found in other individuals of the 1KG cohort. The rest were not found in any of the 1KG individuals and these variants may have low population frequency (<0.1%). Conversely, the 1KG calls for NA12878 contain 224,651 SNVs and 159,163 indels absent from the Platinum catalogue and most of these were observed in our study but were filtered out (see supplementary methods).

### Estimating the extent of platinum genome coverage

To estimate the fraction of the total reference genome that has high-confidence base calls in this study, we combined our Platinum variant catalogue with all high-confidence invariant positions (i.e., explicitly called homozygous reference throughout the pedigree). We concluded that 96.72% of the known reference genome (hg19: autosomes and X chromosome) is covered on the basis of these combined criteria (Table 5) though coverage of the autosomes (97.0%) is significantly better than chromosome X (92.5%) (supplementary methods). Platinum coverage in genes, and particularly exons is higher than the genome average, as expected on the assumption that highly repetitive motifs, which make up most of the non-platinum regions of the reference, are depleted in transcribed and translated sequences compared to the genome average.

**Table 5.** Platinum coverage of the genome, entire genes and exons and in each major category of repeat sequence, based on UCSC (hg19)

| Category | Platinum Bases | Total Bases | % Platinum |
| --- | --- | --- | --- |
| Genome Summary |  |  |  |
| hg19 | 2,742,672,138 | 2,835,673,045 | 96.72 |
| Genes<sup>1</sup> | 1,207,891,444 | 1,240,678,689 | 97.36 |
| Exons<sup>1</sup> | 75,060,473 | 76,594,953 | 98.00 |
| ACMG genes | 4,924,381 | 4,985,610 | 98.77 |
| ACMG exons | 325,531 | 327,215 | 99.49 |
| Repeat Sequence<sup>2</sup> |  |  |  |
| LINE | 586,967,167 | 624,134,772 | 94.04 |
| SINE | 377,522,795 | 389,006,060 | 97.05 |
| LTR | 254,199,704 | 259,386,042 | 98.00 |
| Segmental Duplications | 112,200,846 | 145,135,125 | 77.31 |
| DNA | 96,711,423 | 97,880,579 | 98.81 |
| Simple repeat | 17,832,795 | 25,487,926 | 69.97 |
| Low Complexity | 14,706,936 | 16,652,501 | 88.32 |
| Satellite | 7,990,378 | 11,281,369 | 70.83 |
| Merged Other<sup>3</sup> | 4,904,637 | 6,844,288 | 71.66 |
| All Repeats<sup>4</sup> | 1,417,943,645 | 1,502,863,611 | 94.35 |
| Non Repeats | 1,324,728,493 | 1,332,809,434 | 99.39 |

<sup>1</sup>Breakdown of coverage per gene available in supplemental methods
<sup>2</sup>Certain repeats in the genome may be represented in more than one category
<sup>3</sup>"Merged Other" includes a non-redundant merge of categories listed in RepeatMasker as RNA, rRNA, scRNA, snRNA, srpRNA, tRNA, Unknown and Other.
<sup>4</sup>"All Repeats" is calculated from a single non-redundant merge of all repeat categories.

By this analysis, 93,000,907 reference bases of hg19 (∼3.3%) are classified as non-platinum. However 84,919,966 of these bases lie within segmental duplications or one of the categories of known repeats listed in RepeatMasker, which leaves just 8,080,941 non-platinum bases in the reference that are not accounted for by one of the annotation classes incorporated here. Further work is required to address this remaining 8.1 Mb along with the full characterization of undetected CNVs (see above). This result indicates that we can locate and identify the vast majority of non-platinum bases, that most of them are known repeats, and that they can all be flagged for inclusion or exclusion as dictated by the needs of individual studies in the future.

## Discussion

We combined data from two different sequencing technologies and six different informatics pipelines to identify a ground truth catalogue of 5.4M phased SNVs and indels in this pedigree. A definitive feature of this study is that every platinum variant has been validated using inheritance patterns defined by phased haplotypes in a large pedigree. We avoided the user-defined filters of the mapping and variant calling algorithms, thus maximizing utility of the input data and minimizing potential biases. The Platinum variant set is not intended to be an exhaustive list of variants. Instead, it is a comprehensive genome-wide set of phased small variants that has been validated to high confidence. We discovered that many non-platinum SNVs and small indels in this analysis are likely real, but their inheritance patterns were confounded by co-localised CNVs. For example, the present study highlighted 23,442 locations that are likely to be confounded by CNVs. Next steps will include revision of the assumption of autosomal diploidy so as to include models of multivariate positions. Future work will also need to correct and incorporate the true variants that fail the flanking-sequence analysis; many of these are due to issues with merging and normalizing variant calls from multiple analysis pipelines. A future study might benefit from the use of non-immortalized samples, not confounded by cell culture somatic mutation artifacts. Future studies should also include pedigrees from other ethnic origins.

Compared to other studies, utilizing the genetic inheritance allows us to create a more comprehensive catalogue of platinum variants that reflects both high accuracy and completeness. A comprehensive set of highly-accurate SNVs and indels in both the easy and difficult parts of the genome is vital for software developers because the difficult areas where the current methods most need training data to improve their methods. Additionally, because every one of the variants in this catalogue is phased, this dataset provides a resource to accurately assess emerging technologies designed to provide phasing information (Kitzman et al. 2011; Suk et al. 2011; Zheng et al. 2016).

We demonstrated the use of the platinum variants to make an accurate comparison of different analysis pipelines. This exercise can be repeated using any pipeline to track progress in development and to measure improvements in SNV and indel calling. As tools improve, the same path forward will allow expansion of truth data, with the overriding principle that validation by genetic inheritance enables consideration of variants without being dependent on the strengths and weaknesses of each caller. In particular, 2% of the exons are not within our platinum regions and thus these represent locations where the sequencing workflows fall short. These regions should be prioritised for improvement in future development efforts. In addition, this same method can be used to rapidly assess new genome builds once the inheritance vectors are available and some of the non-platinum variants may be caused by reference issues that will be solved in future genome builds (platinum calls for GRCh38 are also available with the hg19 calls described in this study).

Ultimately a platinum variant catalogue should contain comprehensive, genome-wide sets of all types of variant calls: SNVs, insertions, deletions, large structural rearrangements and CNVs. In this study, a single pedigree provided a very large number of SNVs and indels. Inclusion of CNVs and SVs will become possible as methods for detection improve. However, compared to the small variants, the haplotypes in a single pedigree contain relatively few large structural events. A population-based approach, using aggregated whole genome data, is a more appropriate strategy to identify comprehensive genome-wide sets of CNVs and SVs that can be used to improve large variant callers (Sudmant et al. 2015b). The more common large variants will be identified relatively early on in an aggregate genome study and can then be validated by inheritance in this and other pedigrees. An initial set of common platinum CNVs and SVs will be invaluable to aid algorithm improvement. The rare CNVs and SVs will be discovered over time, as individual genomes are accumulated on a large scale in population-based studies.

## Materials and Methods

### Sequence data and variant calls

Two sets of sequence data from the various members of CEPH pedigree (1463) were generated or used in this study (Table S1). The raw sequence data was aligned to the hg19 reference and SNVs and indels were called using a variety of sequence data and variant calling pipelines (see supplementary methods).

### Calculating inheritance vectors

Inheritance vectors describing the transmission of the parental haplotypes were calculated for the entire genome using the GATK3 SNV calls (DePristo et al. 2011) combined with the linkage software Merlin (Abecasis et al. 2002). Because genotype errors or other deviations from Mendelian inheritance (e.g. hemizygous deletions) will be problematic for the phasing, we only used the SNVs that passed the quality filters in all thirteen samples and showed no Mendelian conflicts. Additionally, some parts of the genome such as the regions around centromeres are prone to mapping errors that may lead to an excess of heterozygous calls that will confound the phasing algorithm. To account for this, we developed an automated approach to merge large regions that show the same familial inheritance but are separated by smaller blocks exhibiting multiple double crossovers. After removing these unlikely crossover events, we were left with 747 distinct inheritance regions for the autosomes and chromosome X. Because we filtered out many SNVs when calculating the initial inheritance vectors, there may be significant gaps where a crossover event occurs. We used the similarity between siblings to narrow these gaps so that the majority of SNVs are included (see supplementary methods). Our final set of defined inheritance blocks cover 2.95 Gb (∼97% of the genome).

### Parsimony analysis

Assuming any variant is biallelic and each sample is diploid, there are a limited number (2^4^−1 = 15 for the autosomes and 2^3^−1=7 for X) of possible phased, genotype combinations in the parents excluding sites that are homozygous reference in every individual. Because the children are not independent of the parents there are also only 15 (7 for X) possible genotype combinations in the entire pedigree within each of the regions defined by the 747 inheritance regions described above. For each of these regions, we calculated the possible genotype combinations across the family defined by the inheritance vectors. For each variant position we compared the observed genotypes against each of the possible genotype combinations based on the known haplotype transmission. If exactly one of the 15 predefined genotype combinations agrees with the observed genotype calls then the site is defined as accurate and by definition phased. See the supplementary methods for a full description of the method and rules used to identify and merge the confident variant calls.

### Data access

Sequence data, merged variant calls and inheritance vectors for the two generation pedigree analyzed here as well as the sequence data for the founders have been submitted to dbGAP (study accepted but waiting on upload instructions). In addition, variant calls described in this work for NA12877 and NA12878 as well as a version of this analysis for GRCh38 are available from www.platinumgenomes.org.

## Acknowledgements

We thank Andrew Gross, Mark Ross and Ryan Taft for helpful discussions and comments.

